# Lesion evidence for a critical role of left posterior but not frontal areas in alpha-beta power decreases during context-driven word production

**DOI:** 10.1101/150748

**Authors:** Vitória Piai, Joost Rommers, Robert T. Knight

## Abstract

Different frequency bands in the electroencephalogram are postulated to support distinct language functions. Studies have suggested that alpha-beta power decreases may index word-retrieval processes. In context-driven word retrieval, participants hear lead-in sentences that either constrain the final word (“He locked the door with the”) or not (“She walked in here with the”). The last word is shown as a picture to be named. Previous studies have consistently found alpha-beta power decreases prior to picture onset for constrained relative to unconstrained sentences, localised to the left lateral-temporal and lateral-frontal lobes. However, the relative contribution of temporal versus frontal areas to alpha-beta power decreases is unknown. We recorded the electroencephalogram from patients with stroke lesions encompassing the left-lateral temporal and inferior parietal regions or left-lateral frontal lobe and from matched controls. Individual-participant analyses revealed a behavioural sentence context facilitation effect in all participants, except for in the two patients with extensive lesions to temporal and inferior-parietal lobes. We replicated the alpha-beta power decreases prior to picture onset in all participants, except for in the two same patients with extensive posterior lesions. Thus, whereas posterior lesions eliminated the behavioural and oscillatory context effect, frontal lesions did not. Hierarchical clustering analyses of the patients’ lesion profiles, and behavioural and electrophysiological effects identified P7 and P9 as having a unique combination of lesion distribution and context effects. These results indicate a critical role for the left lateral-temporal and inferior parietal lobes, but not frontal cortex, in generating the alpha-beta power decreases underlying context-driven word production.

## 1. Introduction

Different frequency bands in the electroencephalogram have been postulated to support distinct language functions (e.g., Lewis, Wang, & Bastiaansen, 2015; McNab, Hillebrand, Swithenby, & Rippon, 2012; Mellem, Bastiaansen, Pilgrim, Medvedev, & Friedman, 2012; Peelle & Davis, 2012; Piai, Roelofs, Rommers, & Maris, 2015). Alpha- and beta-band power decreases in speech production were initially linked to the motor cortex (Salmelin & Sams, 2002; Salmelin, Schnitzler, Schmitz, & Freund, 2000), in line with the motor literature, which has shown a clear relationship between motor preparation and execution and alpha-beta power decreases (e.g., Cheyne, 2013; McFarland, Miner, Vaughan, & Wolpaw, 2000). However, other studies have suggested that alpha-beta oscillations may also index word-retrieval processes, not only in language production but also in language comprehension (Mellem et al., 2012; Piai, Roelofs, & Maris, 2014; Piai et al., 2015; Strauß, Kotz, Scharinger, & Obleser, 2014).

Piai et al. (Piai, Meyer, Dronkers, & Knight, 2017; Piai et al., 2014, 2015) employed a picture-naming task in which the amount of semantic information provided by a preceding sentence was manipulated. Participants named pictures following sentences with a constrained context, such as “He locked the door with the [picture: key]”, or context-neutral sentences, such as “She walked in here with the [picture: key]”. An example of the trial structure is shown in Figure 1A. Picture-naming response times (RTs) are consistently shorter for constrained relative to neutral contexts (Griffin & Bock, 1998; Piai et al., 2014, 2015, 2017). Left-lateralised alpha-beta power decreases for constrained relative to neutral contexts have been consistently found after sentence offset but before picture presentation, in healthy young and older adults (Piai et al., 2014, 2015, 2017). These alpha-beta power decreases have been localised to left angular and supramarginal gyri, left anterior and posterior temporal cortex and left inferior frontal gyrus (LIFG), as shown in Figure 1B (Piai et al., 2015).

**Figure 1.**
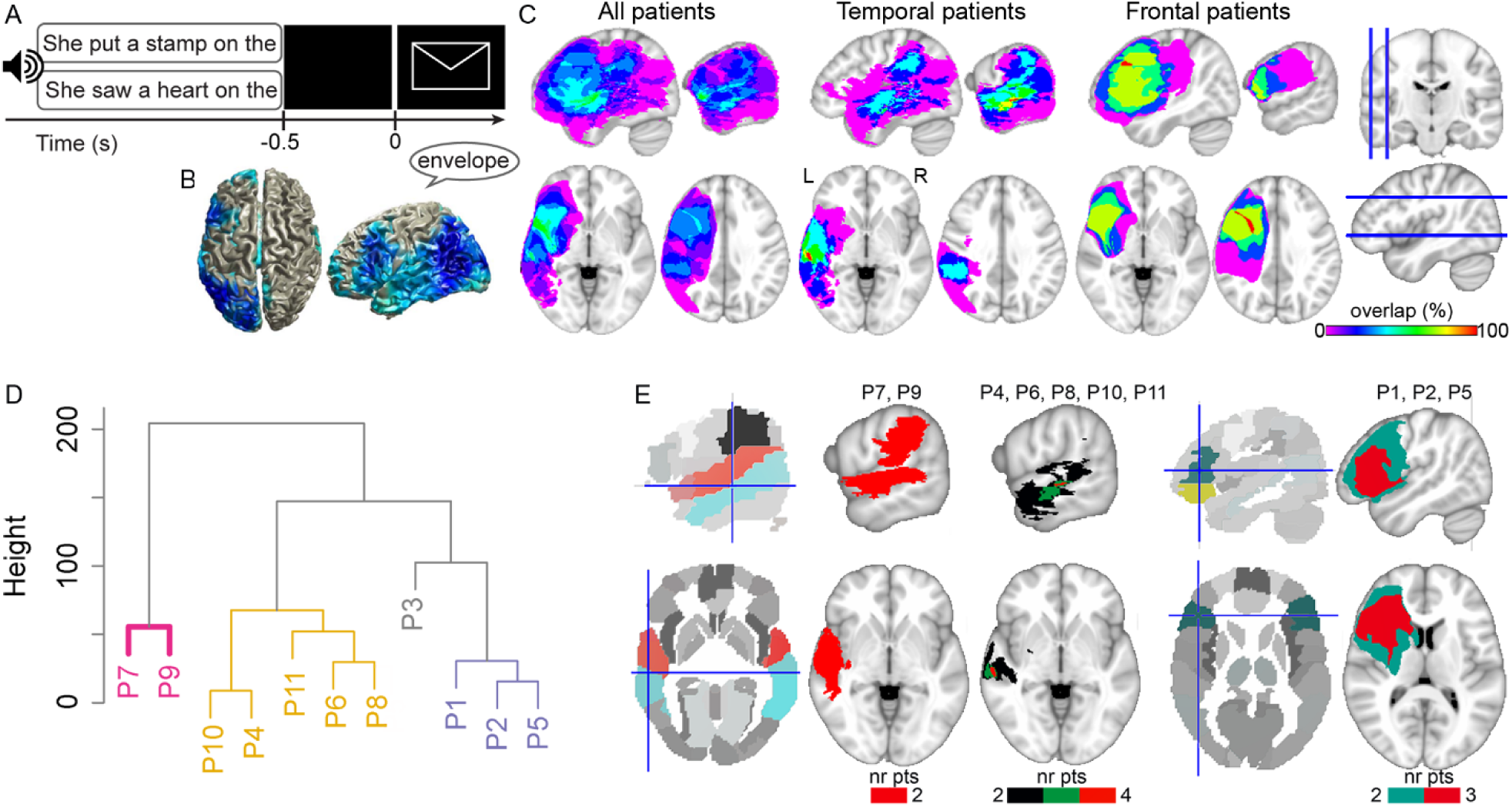
**A.** An example of a trial with constrained (upper) and neutral (lower) sentence contexts with auditory sentences and visual pictures. Only one sentence was presented per trial. **B.** Source localisation of the alpha-beta context effect of Piai et al. (2015). **C.** Lesion overlap map of all patients (left, N = 11), patients with predominantly temporal lobe lesions (middle, N = 6), and patients with predominantly frontal lobe lesions (right, N = 5). The color scale indicates the amount of overlap in lesion location, with magenta indicating that only one patient has a lesion in that particular region (i.e., 0% overlap) and red indicating that all patients have a lesion in that location (i.e., 100% overlap). **D.** Dendrogram of the lesion clusters over left superior temporal, middle temporal, angular, supramarginal, and inferior frontal gyri. Significant clusters are indicated by colours. **E.** Parcellated brains, with relevant regions of interest in colour, and lesion overlap of patients in each cluster (from D), thresholded at where lesions overlap in at least two patients (nr pts = number of patients). Crosshairs indicate left middle temporal gyrus (left) and left inferior frontal gyrus (right).

Given that the scalp-recorded alpha-beta power decreases index a widespread network of sources, it seems unlikely that these power decreases support a single unitary operation. In particular, it is unclear whether some areas are more critical than others for the alpha-beta power decreases to be measurable over the scalp and for behavioural facilitation to occur in the picture naming RTs.

Following previous evidence on the roles of the inferior parietal cortex in conceptual processing (Binder, Desai, Graves, & Conant, 2009) and of the left temporal lobe in lexical retrieval (Baldo, Arévalo, Patterson, & Dronkers, 2013; Indefrey & Levelt, 2004), the alpha-beta power decreases in those areas could reflect conceptual- and lexical-retrieval processes. Evidence for a causal link between alpha-beta power decreases in the left temporal and inferior-parietal lobes and context-driven word production was obtained by Piai et al. (2017). Six patients with left temporal lesions sometimes also including the inferior parietal lobe performed the context-driven picture-naming task. Behavioural facilitation in the picture-naming RTs as well as the alpha-beta power decreases were replicated in four patients. By contrast, two patients with large lesions, encompassing temporal and inferior parietal regions, showed no behavioural facilitation and no alpha-beta power decreases.

These findings help further specify the roles of the left temporal and inferior-parietal lobes to which the sources of the alpha-beta context effect had been localised (Piai et al., 2017), but evidence is lacking on the role of the LIFG in this effect. Using eye tracking, lesion-symptom examinations have suggested a critical role for the LIFG in the integration of information with the ongoing sentence context (Nozari, Mirman, & Thompson-Schill, 2016). However, the relative contribution of left temporal versus left frontal areas to alpha-beta power decreases in word retrieval is unknown.

In the present study, we re-analysed the data of the six patients with lesions to the left temporal cortex (Piai et al., 2017) together with additional data of five patients with lesions to the left frontal cortex (Figure 1C) and ten matched controls. For the controls, we expected to replicate the behavioural context facilitation effect and the alpha-beta power decreases before picture onset (Piai et al., 2014, 2015, 2017). Regarding the patients with left-frontal lesions, based on previous literature indicating the role of the lateral PFC in the use of contextual information to guide behavior (Fogelson, Shah, Scabini, & Knight, 2009; Nozari et al., 2016), we predicted a diminished context effect in the patients whose lesions encompass the lateral prefrontal cortex, and the LIFG in particular.

## 2. Experimental Procedures

The study protocol was approved by the University of California, Berkeley Committee for Protection of Human Subjects, following the declaration of Helsinki. All participants gave written informed consent after the nature of the study was explained and received monetary compensation for their participation.

### 2.1. Participants

Eleven patients with stroke-induced lesions to the left lateral-temporal or lateral-frontal cortex participated (five females; median age = 66, mean = 64, sd = 9, range = 50-74; mean years of education = 17). The distribution of their lesions is shown in Figure 1C. Six patients had lesions predominantly in the left temporal lobe and five in the left frontal lobe. One additional patient with Wernicke’s aphasia and a left temporal lobe lesion was tested. However, almost half (48%) of his responses were errors, so we could not reliably analyse his data. Patients’ language abilities from the Western Aphasia Battery (WAB, Kertesz, 1982) were available for nine patients. Four patients had language abilities within normal limits, according to the WAB. Five patients were classified as anomic, characterised by normal auditory verbal comprehension and repetition, but a relatively impaired word finding ability when speaking. One patient was classified as having conduction aphasia, characterised by normal auditory verbal comprehension, but relatively impaired repetition and word-finding abilities. The two groups of patients did not differ in aphasia severity nor in lesion volume (*t*s < 1, *p*s > .379). All patients were tested at least 12 months post stroke and were pre-morbidly right handed. Information on the patients’ lesions and language ability are shown in Tables 1 and 2.

**Table 1.** Individual lesion volume and percent damage to the left middle temporal gyrus (MTG), superior temporal gyrus (STG), angular gyrus (AG), supramarginal gyrus (SMG), and left inferior frontal gyrus (LIFG).

| Patient | Lesion volume | MTG | STG | AG | SMG | LIFG |
| --- | --- | --- | --- | --- | --- | --- |
| P1 | 52.1 | 0 | 12.9 | 0 | 0 | 59 |
| P2 | 131.76 | 0.1 | 13.1 | 0 | 0 | 93.01 |
| P3 | 122.3 | 0 | 49.8 | 0.5 | 71.6 | 55.1 |
| P4 | 10.09 | 0 | 0 | 0 | 0 | 4.6 |
| P5 | 103.24 | 0 | 10.1 | 0 | 0 | 77.7 |
| P6 | 18.32 | 23.6 | 34 | 2 | 12.5 | 0 |
| P7 | 93.75 | 50.4 | 87.9 | 2.2 | 91.1 | 0 |
| P8 | 103.17 | 17.6 | 33.7 | 5.9 | 32 | 21.4 |
| P9 | 85.82 | 82.6 | 88.6 | 30 | 55.4 | 0 |
| P10 | 4.51 | 6.7 | 3.2 | 0 | 0 | 0 |
| P11 | 36.95 | 56.3 | 22.3 | 0 | 0 | 0 |

**Table 2.**
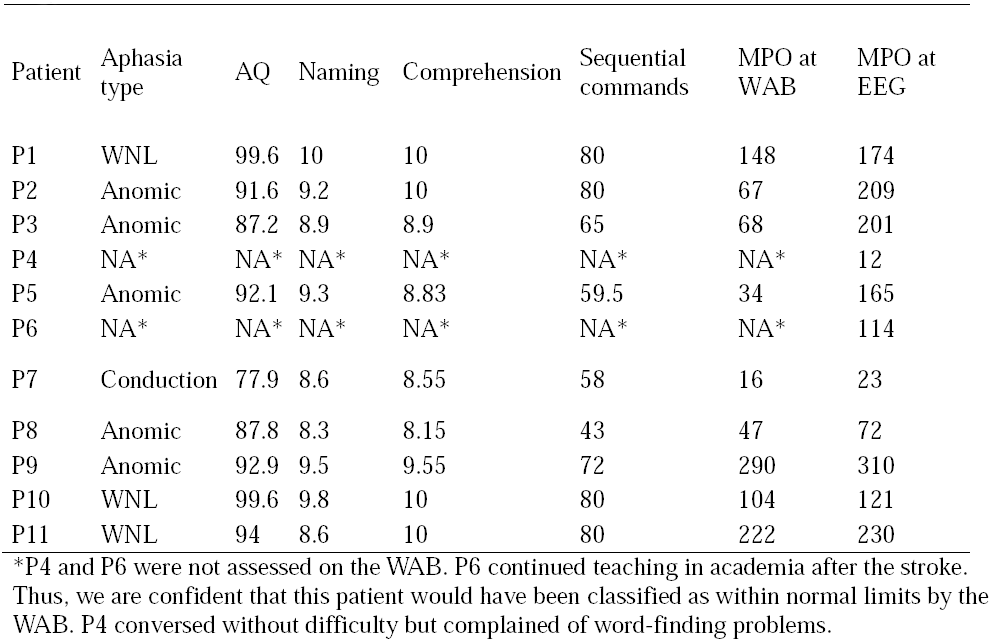
Language testing data from the Western Aphasia Battery (WAB) and time elapsed between stroke date (MPO = months post onset) and WAB testing, and between stroke date and the present EEG experiment. Naming = WAB Naming and Word Finding score (maximum = 10). Comprehension = WAB Auditory Verbal Comprehension score (maximum = 10). Sequential commands = WAB comprehension subtest (maximum = 80). Aphasia Quotient (AQ, maximum = 100). WNL = within normal limit.

Additionally, ten right-handed controls participated, matched for gender, age, and years of education, within ±4 years of age and ±2 years of education to their matched patient (four females; median and median age = 63.5, sd = 8, range = 50-74, *t* < 1, *p* = .838; mean years of education = 17, *t* < 1, *p* > 0.659). None of the patients or control participants had a history of psychiatric disturbances, substance abuse, medical complications, multiple neurological events, or dementia. All participants were native speakers of American English.

### 2.2. Materials

The materials were the same used by Piai et al. (2016) and Piai et al. (2017). Here, we briefly describe the materials but refer the reader to those reports for further detail. Fifty-one colored drawings were selected together with their basic-level name. Each item was paired with two sentences for which the target item completed the sentence. All 102 sentences had six syllables. The sentences belonged to two different conditions. In the neutral condition, no specific word was predictable as the final word of the sentence whereas for the constrained condition, the target word was highly predictable. For each item, the associated sentences had the same two last words. A pre-test confirmed differences in the degree of expectancy for the final word as a function of context (constrained mean cloze probability = 83%; neutral mean cloze probability = 4%, *t*(50) = 45.9, *p* < .001). The sentences in the neutral condition did not have a high cloze probability for any word, with the cloze probability of the most common completion being 23% on average. Sentences were presented auditorily.

### 2.3. Procedure

Stimulus presentation and response recording was controlled by Presentation (Neurobehavioral Systems, Albany, CA). Participants were tested individually in an electrically-shielded, sound-attenuated, dimly-lit booth. In the practice session, participants trained naming the pictures without blinking and postponing their blinking until a cue was given (three asterisks on the screen). The same blinking cue was used during the experiment proper. The sentences were presented via loudspeakers. A trial began with a fixation cross, displayed continuously during auditory sentence playback. After 1 s, the sentence was presented. After sentence offset, the fixation cross remained on the screen for another 0.5 s before the picture was displayed for 2 s. Then, the blinking cue appeared for a variable interval between 1.2 and 1.9 s. An example of an experimental trial is given in Figure 1A.

### 2.4. EEG acquisition

EEG was recorded from 64 Ag/AgCl active scalp electrodes (BIOSEMI, Amsterdam, Netherlands) mounted in an elastic cap according to the extended 10–20 system. EEG was sampled at 1024 Hz. The horizontal electrooculogram was recorded from electrodes placed on the left and right temples and the vertical electrooculogram from Fp1 and the electrode positioned below the left eye. Surface electromyogram was recorded from the orbicularis oris muscle with two electrodes placed on the left upper and right lower corner of the mouth.

### 2.5. Behavioral analysis

Naming responses were monitored online for errors (i.e., disfluent responses, omissions, or incorrect responses). Errors were analysed with logistic regression with condition as a within-participant variable and group as a between-participant variable at an alpha level of 0.05 (two-tailed). Trials corresponding to errors were subsequently excluded from all RT and EEG analyses. Response times were calculated manually using Praat (Boersma & Weenink, 2013) before the trials were separated by condition. Statistical analyses of the RTs were conducted using R (R Development Core Team, 2014).

Participants’ median RTs were computed and an analysis of variance was run on the RTs, with context condition (constrained vs neutral) as a within-participant variable and group (controls vs patients) as a between-participant variable at an alpha level of .05 (two-tailed). The context effect was also examined at the individual-participant level using independent-samples *t*-test across trials.

### 2.6. Lesion analysis

Lesions were drawn on patients’ structural magnetic resonance images (MRIs) by a trained technician and confirmed by a neurologist (RTK). Lesions masks were then normalised to the MNI template. Percent damage to different areas were determined based on the Automated Anatomical Labeling template in MRIcroN (Rorden, Karnath, & Bonilha, 2007). To investigate the patients’ lesion profile, we used hierarchical clustering over the percentage of damage of the left middle temporal gyrus (MTG), superior temporal gyrus (STG), angular gyrus (AG), supramarginal gyrus (SMG), and LIFG (all entered as separate variables). Cluster analysis is a form of unsupervised learning, used to find structure in the data. It is employed to group elements in so-called clusters such that elements in one same cluster are more similar to each other than to elements in other clusters. The Euclidean distance was used as the distance measure between pairs of observations and the Ward’s criterion was used as the linkage criterion. To validate the cluster solution, multiscale bootstrap resampling was employed with 5,000 bootstraps (Suzuki & Shimodaira, 2006). *P* values, indicating how well the clusters are supported by the data, were derived from the Approximately Unbiased *p* value (Suzuki & Shimodaira, 2006) and only clusters below an alpha level of .05 are reported.

### 2.7. EEG analysis

The analyses were performed using FieldTrip version 20160619 (Oostenveld, Fries, Maris, & Schoffelen, 2011) in MatlabR2014a. Each electrode was re-referenced off-line to averaged mastoids. The data were high-pass filtered at 0.16 Hz (FieldTrip default filter settings) and segmented into epochs time-locked to picture presentation, from 800 ms pre-picture onset to 300 ms post-picture onset. All epochs were inspected individually for artifacts such as eye-movements, blinks, muscle activity, and electrode drifting. For two participants (P6 and C14), rejecting trials with eye blinks resulted in substantial data loss. Therefore, for these two participants, independent component analysis was used to correct for blinks (Jung et al., 2000, as implemented in FieldTrip). Eight peripheral channels (T7, T8, TP7, TP8, F7, F8, FT7, FT8) were excessively noisy (variance > 4 millivolt) in the majority of participants and were removed from analyses. On average, error- and artifact-free trials comprised 46 trials per condition for controls (no difference in trial numbers between conditions, *t*(9) < 1, *p* = .435) and 44 trials for patients (no difference in trial numbers between conditions, *t*(10) < 1, *p* = .910). The number of available trials for patients was on average only two fewer than for controls, *t*(19) = -1.795, *p* = .089). Time-resolved spectra were calculated with a modified spectrogram approach, at frequencies ranging from 8 to 25 Hz (following findings from Piai et al. 2014, 2015, 2017), with an adaptive sliding time window of three cycles’ length (e.g., the window was 300 ms long at 10 Hz). This window was advanced in steps of 10 ms in the time dimension and in steps of 1 Hz in the frequency dimension. The data in each window was multiplied with a Hanning taper, and the Fourier transform was taken from the tapered signal. Note that no baseline correction was used.

#### 2.7.1. Statistical analysis

For each group (patients and controls), we compared the time-frequency representations between the two conditions using a non-parametric cluster-based permutation test (Maris & Oostenveld, 2007). The test was performed on all available channels, time points (i.e., -800 ms to 300 ms relative to picture onset), and frequencies (i.e., 8-25 Hz). This three-dimensional space was first scanned for time points, frequencies, and channels that exhibited a similar difference between the two conditions across participants based on a two-tailed dependent-samples t-tests at an alpha level of .05. Time points, frequencies, and channels whose *p* values are lower than the alpha level are selected and clustered on the basis of adjacency. For each cluster, a cluster-level statistic was then calculated by taking the sum of the *t*-values within that cluster. The clusters’ statistical significance was then calculated with a Monte Carlo method, for which a permutation distribution is created by randomly partitioning the data into two conditions. Then, the same scanning and clustering procedure is performed for each random partition and the cluster with the largest summed *t*-values is selected to enter the permutation distribution. This procedure is repeated 1,000 times. All cluster-level statistics from the observed data are then compared to this permutation distribution and the proportion of random partitions that yielded a larger cluster test-statistic than that of the observed cluster represents the Monte Carlo estimate of the *p* value. Using a critical alpha-level of .05, we conclude that the constrained and neutral conditions differ from each other significantly if this Monte Carlo *p*-value is smaller than .05.

We also assessed group differences by comparing the relative power differences between the two conditions (i.e., power differences divided by the averaged power across conditions) as a function of group. The same cluster-based permutation procedure was adopted, with the difference that independent-samples *t* tests were used.

### 2.8. Clustering of variables

A hierarchical clustering approach was used to divide patients into groups as a function of the anatomical and functional variables available. These variables were the following: 1) the percentage of damage of the left MTG, left STG, left AG, left SMG, and LIFG (as explained above), 2) lesion volume, 3) the behavioural context effect (the mean difference between the two conditions), and 4) the EEG context effect, that is, the relative power differences between the two conditions averaged over all available channels in the 8-25 Hz range between -300 ms and picture onset (i.e., the significant cluster on the group level, see Results below). The same procedure for the clustering analysis was used as reported above (“Lesion analysis”), except that clusters were evaluated at an alpha level of .0125.

## 3. Results

### 3.1. Lesion profile

Figure 1D shows how patients are grouped as a function of their lesion profile. The y-axis indicates how dissimilar, according to the Euclidean distance, the individual data points and clusters are from each other. Significant clusters are indicated by colours. As can be seen in Figure 1D, patients P7 and P9 are grouped together and separately from the other patients. This means that the lesion profile of these two patients is more similar to each other than to the lesion profile of the patients in other clusters. The same is the case for the other clusters. The lesion overlap of the patients in each cluster is shown in Figure 1E thresholded at where lesions overlap in at least two patients. Patients P7 and P9 have a lesion in STG, MTG, and SMG. Patients P4, P6, P8, P10, and P11 form a second cluster, with no 100% lesion overlap among them, but with four patients having an overlapping lesion in MTG. Finally, patients P1, P2, and P5 form a third significant cluster, with all three patients in this cluster having a predominantly LIFG (and insular) lesion.

### 3.2. Sentence context facilitates picture naming

Table 3 shows the error rates for each patient per condition. Control participants made fewer errors than patients (group-level: 0.78% vs 5% respectively, beta = -2.387, S.E. = .749, z-value = -3.226, p < .002). The remaining comparisons were not statistically significant, all *p*s > .176.

**Table 3.**
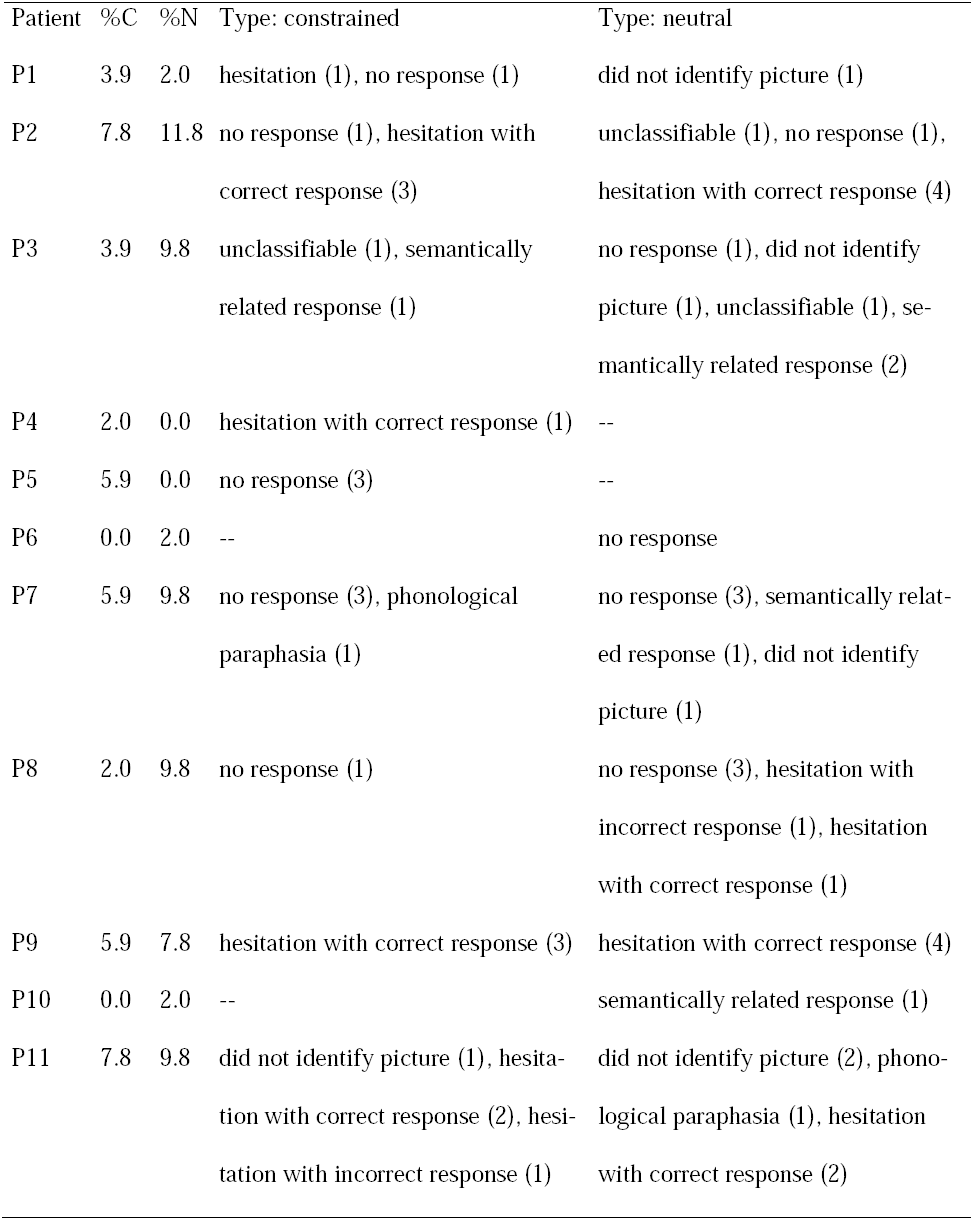
Error rate (in percentage) and error type per patient and condition. C = constrained; N = neutral.

| Patient | %C | %N | Type: constrained | Type: neutral |
| --- | --- | --- | --- | --- |
| P1 | 3.9 | 2.0 | hesitation (1), no response (1) | did not identify picture (1) |
| P2 | 7.8 | 11.8 | no response (1), hesitation with correct response (3) | unclassifiable (1), no response (1), hesitation with correct response (4) |
| P3 | 3.9 | 9.8 | unclassifiable (1), semantically related response (1) | no response (1), did not identify picture (1), unclassifiable (1), semantically related response (2) |
| P4 | 2.0 | 0.0 | hesitation with correct response (1) | -- |
| P5 | 5.9 | 0.0 | no response (3) | -- |
| P6 | 0.0 | 2.0 | -- | no response |
| P7 | 5.9 | 9.8 | no response (3), phonological paraphasia (1) | no response (3), semantically related response (1), did not identify picture (1) |
| P8 | 2.0 | 9.8 | no response (1) | no response (3), hesitation with incorrect response (1), hesitation with correct response (1) |
| P9 | 5.9 | 7.8 | hesitation with correct response (3) | hesitation with correct response (4) |
| P10 | 0.0 | 2.0 | -- | semantically related response (1) |
| P11 | 7.8 | 9.8 | did not identify picture (1), hesitation with correct response (2), hesitation with incorrect response (1) | did not identify picture (2), phonological paraphasia (1), hesitation with correct response (2) |

Figure 2 shows the participant-level median RTs per condition (left panel) and the individual magnitude and significance of the context effect, with 95% confidence intervals (right panel). On the group level, responses in the constrained condition were faster than in the neutral condition (*F*(1,19) = 101, *p* < .001) and patients were overall slower than controls (*F*(1,19) = 5.05, *p* = .037), but the interaction was not significant (*F*(1,19) < 1). On the individual level, all control participants and all but two patients (P7 and P9) were reliably faster on constrained than neutral trials, indicating a robust within-participant behavioral facilitation effect.

**Figure 2.**
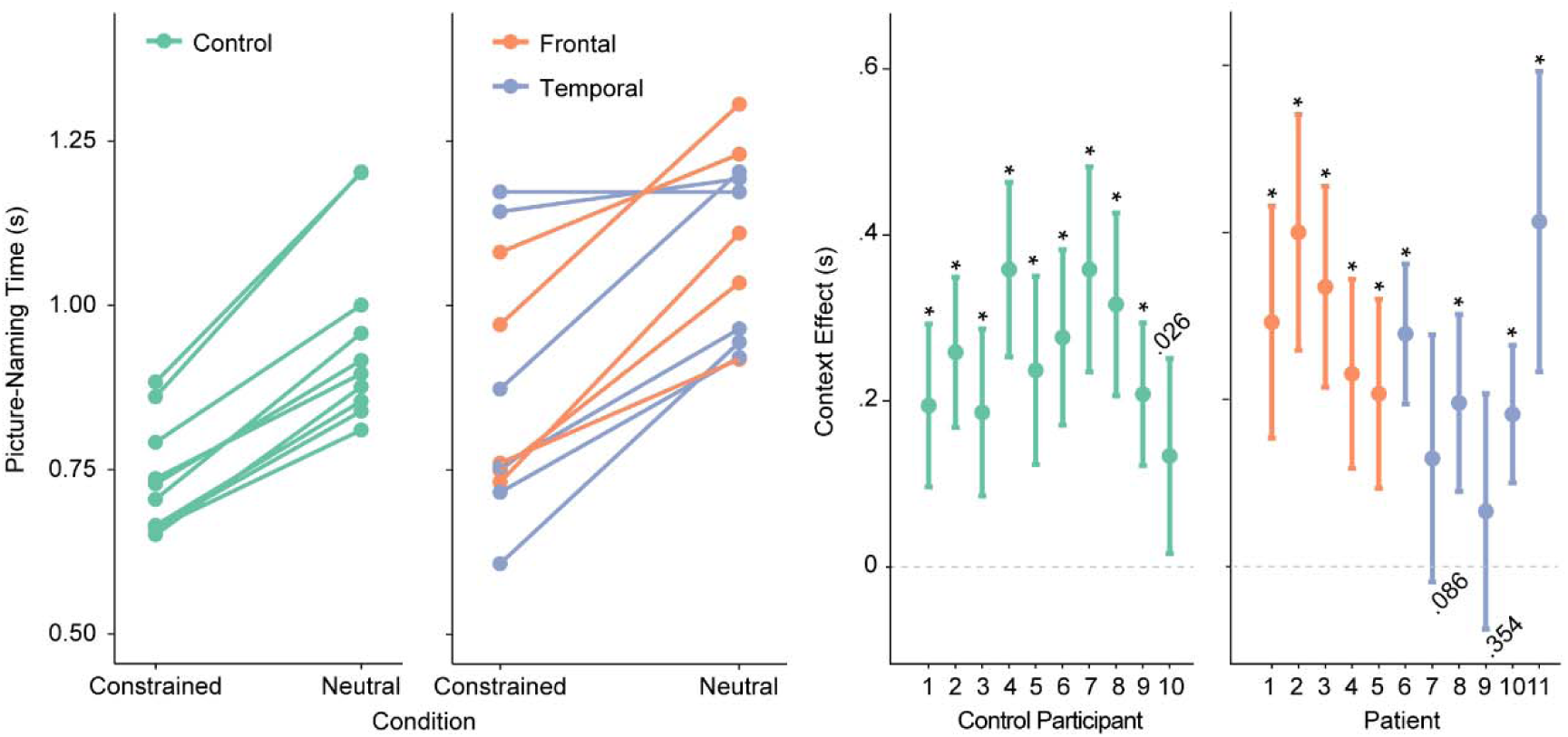
**A.** Median picture naming times for each participant and condition. **B.** Context facilitation effect (neutral – constrained) with 95% confidence intervals and *p* value for each participant. Asterisks indicate *p* < .001.

### 3.3. Sentence context modulates pre-picture alpha-beta power

In both controls and patients, we observed 15%-20% 15-20% power decreases (relative to the average across conditions) in the alpha-beta range prior to picture presentation, as shown in Figure 3A. Cluster-based permutation tests indicated a statistically significant context effect for both controls (Monte Carlo *p =* .039) and patients (Monte Carlo *p <* .001) that could be attributed to a spatio-spectro-temporal cluster of power decreases in each group. These clusters were detected between 8 and 25 Hz and -0.3 to 0 s in both groups, in the channels indicated in white in the topographies. The interaction between context condition and group did not yield any significant clusters (both when all patients were included and when patients P7 and P9 were excluded). This result is to be expected given that, on the group level, both groups show a significant context effect associated with a cluster in the 8-25 Hz range prior to picture onset.

**Figure 3.**
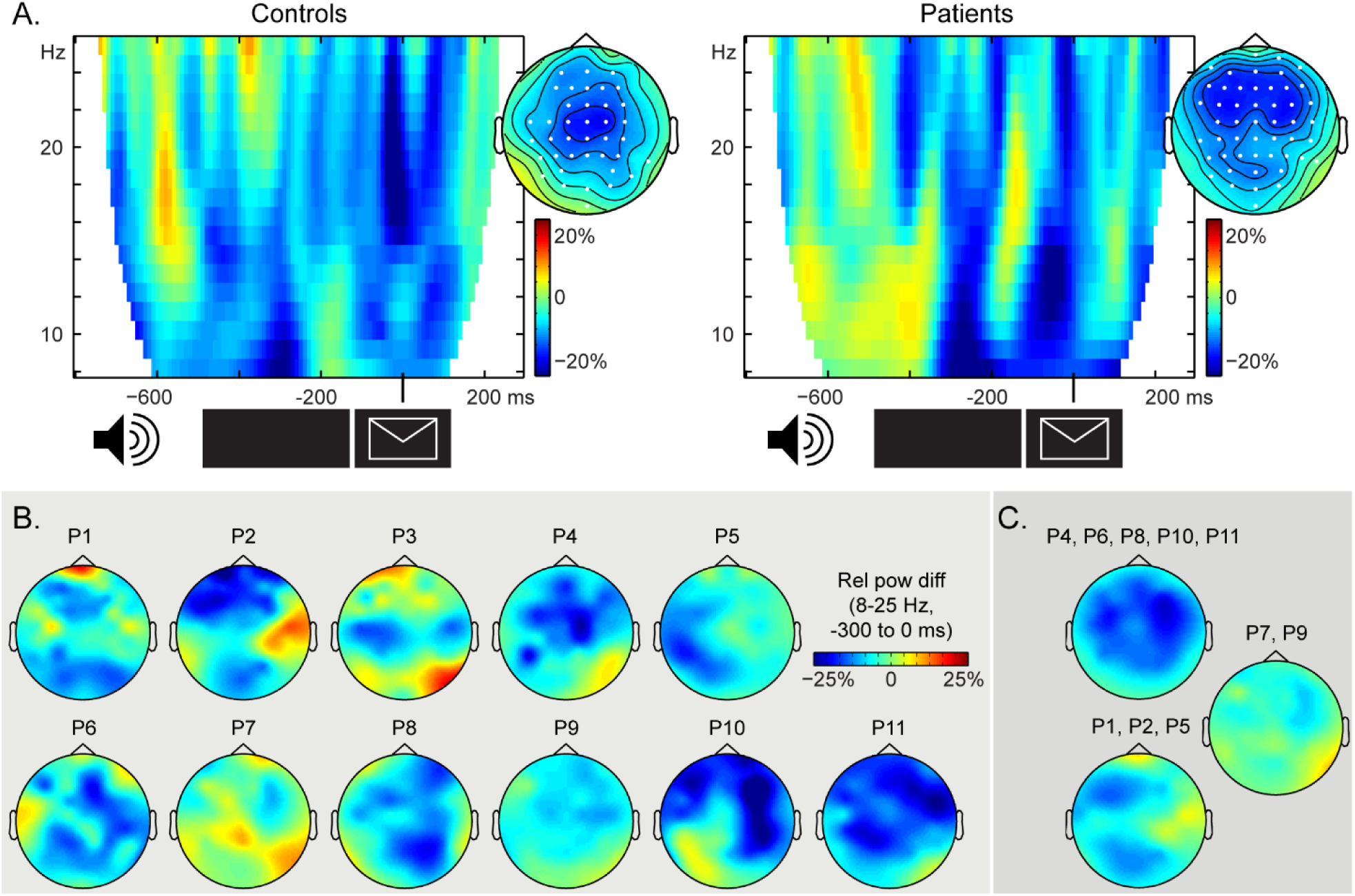
**A.** Time-frequency representations of the context effect (in relative power changes) averaged over the channels associated with the significant clusters, indicated in white on the topographical maps. The topographical distributions of the significant cluster (8 to 25 Hz, -0.3 to 0 s) are shown for each group. **B.** Patients’ individual topographical distribution of the context effect (8 to 25 Hz, -0.3 to 0 s), expressed as relative power changes. **C.** Topographical distribution of the context effect (in relative power changes) averaged over the patients in each cluster from Figure 1D.

Individual EEG effects are shown in Figure 3B. Patients P7 and P9 showed the weakest alpha-beta power decreases over the scalp. EEG effects averaged over clusters of patients based on their lesion profiles (Figure 1D) are shown in Figure 3C. These topographies indicate that for the patients with lesions overlapping in the mid portion of the MTG (P4, P6, P8, P10, P11, the yellow cluster in Figure 1D), the scalp effect is bilateral, with a slight right-lateralised bias. For the patients with a predominantly LIFG lesion (P1, P2, P5, the purple cluster in Figure 1D), the scalp effect is largely left-lateralised. The average over P7 and P9 (the pink in Figure 1D) confirms that these two patients have the weakest EEG effects.

### 3.4. Extensive posterior lesions and weak behavioural and oscillatory effects co-occur

Figure 4 shows the results of the hierarchical clustering analyses. Significant clusters are indicated by filled lines. When all functional and anatomical variables were entered, the clustering procedure yielded one distinct significant cluster, with patients P7 and P9 separately from the other patients. We ran additional analyses to further examine the uniqueness of this clustering solution. When clustering was performed on all variables but the EEG (i.e., pre-picture alpha-beta power), patients P7 and P9 were again clustered together, separately from the other patients. Performing the clustering procedure on all variables but lesion volume also yielded a unique cluster of patients P7 and P9. Finally, when all variables were entered but the behavioural effect, patients P7 and P9 again clustered together. In summary, despite the changing configuration of the dendrograms with different variables entered in the clustering analysis, patients P7 and P9 had a unique constellation of functional and anatomical variables different from the other patients.

**Figure 4.**
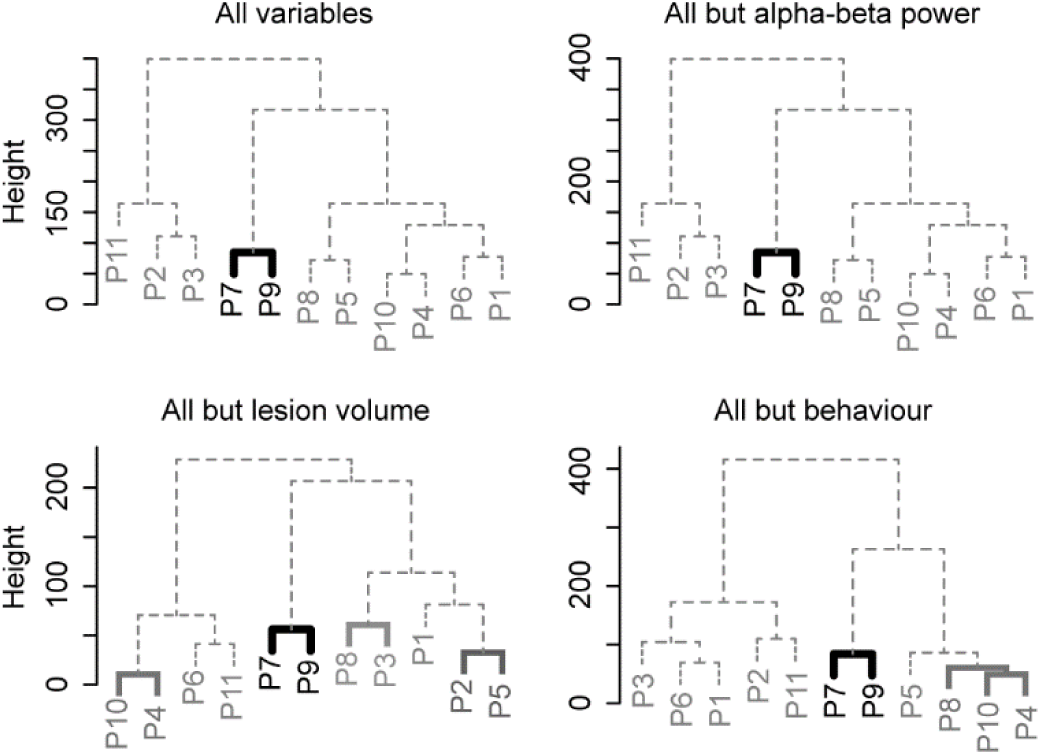
Dendrograms of the patient clusters for all functional (EEG, behaviour) and anatomical (percent lesion in regions of interest, lesion volume) variables (top left), all variables but EEG (top right), all variables but lesion volume (bottom left), and all variables but behaviour (bottom right). Significant clusters (at an alpha level of .0125) are indicated by colours.

## 4. Discussion

In the present study, patients with stroke-induced lesions to the left temporal, inferior parietal, or frontal lobes and matched controls performed a context-driven picture naming task while their EEG was recorded.

Replicating previous findings (Griffin & Bock, 1998; Piai et al., 2014, 2015, 2017), all healthy-control participants showed shorter picture naming times and alpha-beta power decreases before picture onset following constrained sentences relative to neutral sentences. This pattern of behavioural and electrophysiological effects was also found for all patients, except for two patients with extensive lesions to the temporal and inferior parietal lobes (P7 and P9). We used hierarchical clustering to group patients as a function of their lesions profiles, and behavioural and oscillatory effects. These analyses indicated that P7 and P9 had a unique combination of lesion distribution and lack of behavioural and electrophysiological context effects. By contrast, no such association was found between the LIFG and alpha-beta power decreases. Together, these results provide evidence for a causal link between posterior alpha-beta power decreases (i.e., in the left lateral-temporal and inferior parietal lobes) and context-driven word retrieval.

Regarding the naming latencies, the fact that patients with PFC lesions still benefitted from constraining sentence contexts is noteworthy as previous studies have highlighted the critical role of the lateral PFC in people’s ability to use contextual information to guide behavior. Fogelson et al. (2009) examined seven patients with lateral PFC lesions (with the greatest lesion overlap in the dorsolateral PFC) performing a non-verbal target-detection task in which patients responded manually to a target following a predictive or a random sequence. Both behavioural and neurophysiological evidence was obtained for patients’ impaired ability to employ the local context (i.e., the sequence) to anticipate the target and respond to it, contrary to our findings. However, the predictive context in their study consisted of abstract visual sequences, whereas in our study, the sequences consisted of meaningful words. This aspect of our materials may have decreased the demands on control and selection processes, supported by the lateral PFC.

A similar account is likely for the differences between our findings and those of Nozari et al. (2016). Using eye tracking, Nozari et al. compared the anticipatory eye-movements of four patients with LIFG lesions to those of three patients with lesions to the left temporal and parietal lobes. Patients watched a screen with four pictures (e.g., car, hat, banana, flashlight) while hearing sentences with restrictive or non-restrictive verbs (e.g., “She will drive the” vs “She will study the”). The patients with left posterior lesions looked at the target (here, the pictured car) earlier following a restrictive verb than the patients with LIFG lesions. The authors concluded that LIFG damage impaired patients’ ability to use the semantic contextual cues present in the sentence to relate it to the target picture. However, the visual-world paradigm employed by Nozari et al. (2015) requires participants to integrate visual information from the pictures with the auditory sentence and to avoid influences from irrelevant pictures in the display, possibly increasing control and selection demands on the PFC. In our case, spreading activation in the lexico-semantic network through the information in the sentence alone may have been sufficient to guide retrieval of the picture concept and its name.

The present findings begin to illuminate the relative roles of left temporal, inferior parietal, and frontal areas in the alpha-beta context effect in word production. Whereas posterior lesions, including the left temporal and inferior parietal lobes, eliminated alpha-beta power decreases prior to picture onset, LIFG lesions did not. Thus, out of the various sources implicated in eliciting the context alpha-beta power decreases (Piai et al., 2015), the temporal and inferior parietal areas might play a more critical role than the left inferior frontal areas. Given the functional roles associated with these brain areas, alpha-beta power decreases could reflect a mix of core semantic memory and lexical retrieval processes (Baldo et al., 2013; Binder et al., 2009; Schwartz et al., 2009), along with additional frontal control processes that may have been less critical in the task employed in this study.

Previous studies have found correlational evidence for the role of alpha-beta power decreases in lexical retrieval (e.g., Brennan, Lignos, Embick, & Roberts, 2014; Mellem et al., 2012; Piai et al., 2015). The present study contributes a causal link between alpha-beta power decreases in the left temporal and inferior-parietal lobes and context-driven word production. However, it is still unclear whether one sole oscillatory signature (i.e., alpha-beta power decreases) indexes lexical retrieval or whether distinct aspects of this process are indexed by different frequency bands (e.g., Bastiaansen, van der Linden, Ter Keurs, Dijkstra, & Hagoort, 2005; Marinković, Rosen, Cox, & Kovacevic, 2012; Mellem et al., 2012; Piai et al., 2014). Future studies will hopefully clarify this issue.

Understanding language-related alpha-beta oscillations is not only relevant for theory but also for improving clinical applications of these neuronal signatures. Previous studies have used alpha-beta power modulations as an index of language function to understand hemispheric functional re-organisation after left-hemisphere strokes (Kielar, Deschamps, Jokel, & Meltzer, 2016; Meltzer, Wagage, Ryder, Solomon, & Braun, 2013; Piai et al., 2017) and to determine hemispheric dominance for language preoperatively in neurosurgical patients (Findlay et al., 2012). We suggest that theoretical and clinical progress on this subject will be mutually informative to both the neurobiology of language and patient care.

In conclusion, we obtained behavioural and neurophysiological evidence for a causal link between alpha-beta power decreases in the left lateral-temporal and inferior parietal lobes and context-driven word production.

## Acknowledgements

The authors are grateful for the patients and their families, as well as for the other volunteer participants for taking part in this study. We would like to thank Donatella Scabini and Brian Curran for patient delineation, Brian Curran, Clay Clayworth and Callum Dewar for lesion reconstruction, Amber Moncrief and Selvi Paulraj for helping design the materials, Paige Mumford and Laura Agee for help with audio recordings, Kristoffer Dahlslätt for invaluable discussions, and the members of the Center for Aphasia and Related Disorders at the VAHCS in Martinez, CA, for neuropsychological testing. The authors declare no conflict of interest.

### Funding

This work is supported by grants from the Netherlands Organization for Scientific Research (446-13-009 to V.P., 275-89-032 to J.R.), the National Institutes of Health (NINDS R37 NS21135 to R.T.K), and by the Nielsen Corporation.

## Data sharing statement

This article’s supporting data and materials can be obtained upon request.

## Author contributions

Conception and design of the work: VP

Data collection: VP

Data analysis and interpretation: VP

Drafting the article: VP, JR

Critical revision of the article: VP, JR, RTK

Final approval of the version to be published: VP, JR, RTK

